# Multi-task Learning and Ensemble Approach to Predict Cognitive Scores for Patients with Alzheimer’s Disease

**DOI:** 10.1101/2021.12.08.471856

**Authors:** Daren Ma, Christabelle Pabalan, Abhejit Rajagopal, Akanksha Akanksha, Yannet Interian, Yang Yang, Ashish Raj

## Abstract

During its chronic degenerative course, Alzheimer’s Disease severely harms the patients’ cognitive abilities. Assessment of current and future cognition is an integral component of a diagnosis of dementia, and therefore an important clinical and scientific goal. Unfortunately, subjective, time-consuming and operator-sensitive clinical surveys or neuropyschiatric batteries remain the only viable methods of assessing cognition. Given that MRI is the most prevalent, cost-effective, and clinically important imaging modality, it may be considered a suitable predictor of cognition. Yet, it has hitherto proved very challenging to predict one from the other. We propose that an image-based Deep Learning model can be custom-built to achieve this goal. We designed a novel multi-task UNet model to predict the subjects’ current and future cognition (via ADAS-Cog scores), taking as input baseline T1-weighted MRI and demographic risk factors. The key innovation in the model is that it seeks to solve two adjacent but relevant tasks: image segmentation into tissue types; and prediction of cognition. The first task gives a high-accuracy brain segmentation, comparable to other cutting edge methods. The features trained from the segmentation task are used in the cognition task. This combination is far superior to stand-alone single-shot cognition models. We achieved excellent accuracy in both baseline and time-series forecast of ADAS-Cog scores. Through further feature map analysis made on the receptive fields, we managed to impart much-needed model interpretability, critical for real-world clinical practice. This study constitutes the best-reported performance of any comparable approach, and opens the door towards machine-based tracking of AD progression.

## 1. Introduction

Alzheimer’s disease (AD) is a neurodegenerative disorder affecting upwards of 50 million people and 60-70% of dementia cases worldwide. Clinically, AD is diagnosed based on history of the illness, cognitive tests, and by ruling out other causes of symptoms. Many risk factors of its progression are known, including apolipoproteins (APO), Angiotensin-converting enzyme (ACE) inhibitors, amyloid plaques, neurofibrillary tangles, and loss of neural pathways Farrer et al. (2000); Shoghi-Jadid et al. (2002); Genin et al. (2011). While postmortem histopathology remains the only gold standard, there has been persistent interest to quantify progression using non-invasive magnetic resonance imaging (MRI) Frisoni et al. (2010); Nordberg et al. (2010). Therefore there is tremendous interest in using modern machine learning (ML) methods on MRI for early detection, diagnosis and prognosis, with recent focus on deep learning (DL) techniques based on deep neural networks (DNNs) – we include a brief survey in Discussion. These approaches have primarily focused on disease diagnosis and on predicting conversion to dementia - essentially classification tasks.

ML methods for regression tasks using neuroimaging are less frequent and far less successful Sui et al. (2020). Arguably, binary diagnostic classification is frequently less useful clinically than predicting a patient’s cognitive status, which gives a clinically meaningful benchmark of disease severity and its practical impact in the patient’s life. Disease severity and progression rates are themselves closely associated with cognitive score Ito et al. (2011), hence an accurate prediction of cognition will also indirectly aid diagnosis and prognosis. Many clinical measures of cognitive performance are currently available; they typically utilize a battery of neuropsychiatric and neurocognitive tests delivered by a trained neuropsychologist. Of these, the Alzheimer’s Disease Assessment Scale Cognitive (ADAS-Cog) score is one of the most important standards Rozzini et al. (2007).

In this work we address the task of predicting a patient’s current and future ADAS-Cog trajectory from their baseline MRI and demographic information. This has utility in early diagnosis and prognosis, but may also inform new clinical protocols. For example, incorporating cognitive scores and MRI biomarkers could yield accurate prediction ahead of dementia symptoms Zandifar et al. (2020), which may help to customize therapies and lead to early effective treatments.

Deep learning would appear to be an excellent tool for predicting ADAS-Cog, since it can remove the reliance on time-consuming preprocessing and prior feature selection, and thus avoiding the related model-dependent decisions (Sui et al., 2020). However, the success of ML/DL methods on classification tasks has not translated easily to prediction of cognitive scores Sui et al. (2020). We show that this problem is not effectively solved using existing off-the-shelf ML/DL tools. A conventional single-shot CNN tasked with this problem suffers from the extreme dimensionality mismatch between voxel-level imaging input and scalar cognitive output, hence is and severely hampered by a lack of sufficiently large sample sizes.

Instead, we introduce a bespoke, hybrid DL approach that combines demographic data and dense features extracted by a 3D UNet as input to a gradient boosting algorithm that regresses directly on the ADAS-Cog11 score. We propose that the cognitive task is made more effective by solving an adjacent but relevant low-level image task - that of image segmentation. We use MRI rather than molecular PET imaging (which typically contains pathology information) because MRI remains the most prevalent, cost-effective, and clinically utilized neuroimaging modality with widespread community access. Further, since the brain’s cognitive performance is mediated (albeit in a complex manner) by its morphology and organization, the excellent soft-tissue contrast of MRI should make it a good predictor of cognition.

To capture the cognition-sensitive structure, our approach uses features derived from the fully-encoded (lowest dimensional) layer of a 3D residual UNet trained on the tissue segmentation task. This design ensures that training for the segmentation task produces model features that will also be relevant for cognition, and results in implicitly-conditioned neuroimaging features. While the segmentation task serves as a regularization tool and to enable efficient learning in a data-scarce setting, we note its importance in its own right for clinicians and neuroimaging scientists. Traditionally, the segmentation task was performed using dedicated advanced and computationally-expensive neuroimaging pipelines such as FreeSurfer Fischl (2012). The ability to achieve this output within seconds is in itself a valuable secondary product. Further, as we shall see, the reuse of the segmentation maps’ tissue volumes is critical to correctly predict longitudinal changes in cognition.

Our ML pipeline has three phases: a UNet segmentation model to segment MRI scans; a multi-task model that additionally performs a regression task on cognitive scores; and an ensemble model that performs both segmentation and regression; see outline in Figure 1. We show that this multitask framework is indeed highly effective at both its tasks, producing high-quality segmentation maps of the brain as well as highly accurate cognitive scores. For comparison we also implemented the base CNN model without the bespoke multitask design – it gave significantly poorer performance. Extending this approach to longitudinal data and re-utilising the volumetric output produced by the segmentation task, our model gives promising predictions of cognition up to 48 months out using baseline images. Our model easily outperforms other recent methods (Section 5.3). We are aware of no other equally successful predictive model of future cognitive scores from baseline imaging.

**Fig. 1.**
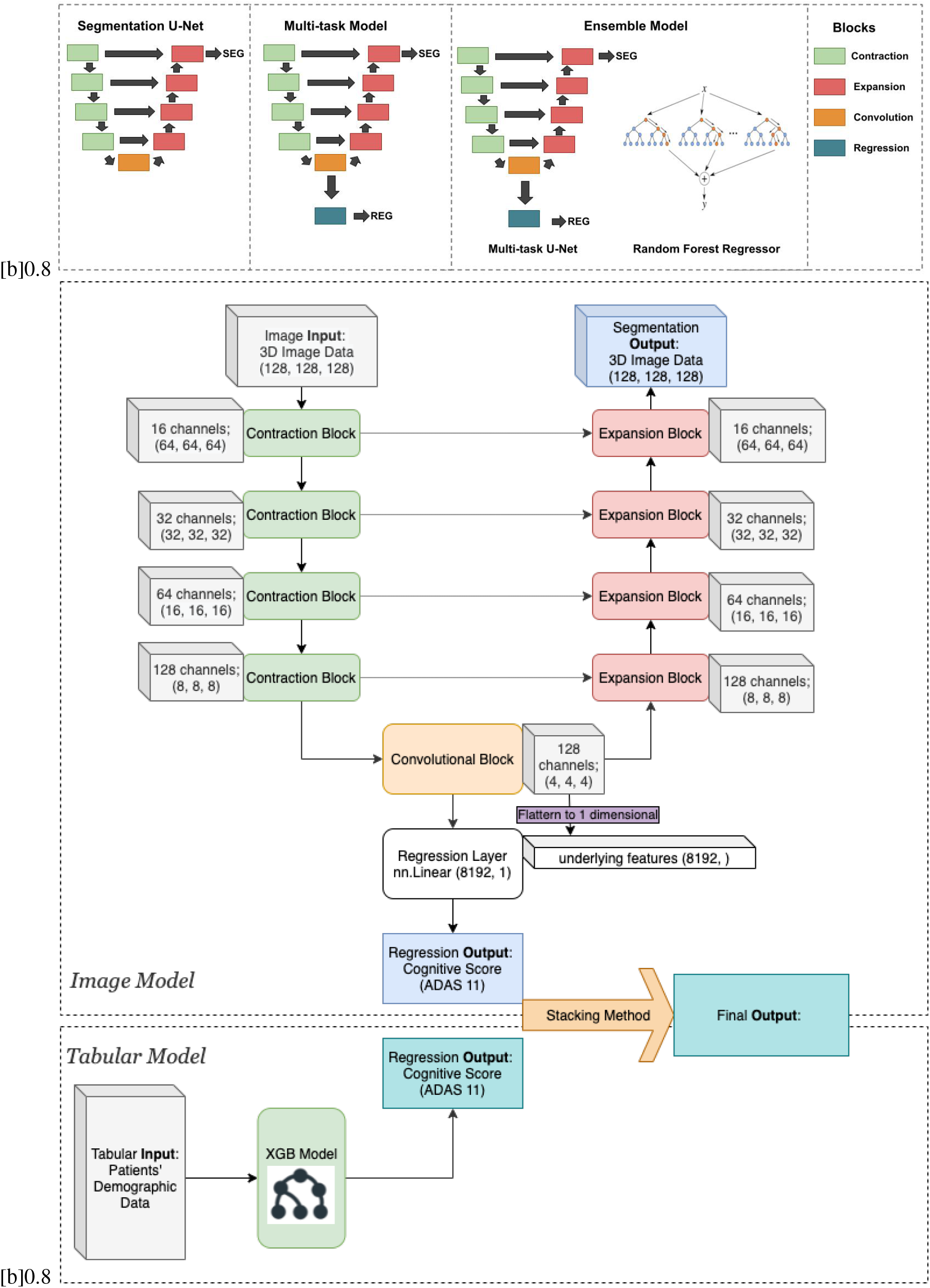
Overall model architecture. Top: Incrementally embellished development of the model, with the basic segmentation 3D UNet augmented by the regression layer for cognition, and then by the non-imaging tree-based regression model. Middle: Detailed structure of the custom multitask UNet model is trained on the tissue segmentation task, and the features from its most concentrated convolutional block is taken for the purpose of predicting cognitive scores, after flattening the 128 channels of 4x4x4 matrices into a feature vector of length 8192. Bottom: Random Forest regression model fits the demographic data to cognition. These predictors are stacked in an ensemble model to generate a better estimate of ADAS-Cog11.

## 2. Methods

In this work, a multitask model will be created, such that the subjects’ ADAS-Cog11 scores are our major target, and the MRI segmentation as our supportive target.

### 2.1. Dataset

The major training data were obtained from the Alzheimer’s Disease Neuroimaging Initiative (ADNI) database (http://adni.loni.usc.edu), which contains longitudinal cognitive, demographic, fluid, genetic and imaging data on a large sample of AD and control subjects. The study was approved by the Institutional Review Boards at each ADNI site. Informed consent was obtained from all subjects prior to enrollment. All methods were carried out in accordance with relevant guidelines and regulations.

For this study we found 2288 subjects in the ADNI 1, 2, 3, and GO metadata dataframe, of which 1952 subjects with MRI and diagnosis were available. From these we selected 1839 subjects, who had (a) current diagnosis status of AD, MCI or Control; and (b) a valid MRI image record for at least one visit. To enlarge the training samples for the segmentation task, we also gathered 1181 healthy youth brain MRIs from HCP, age ranging from 20 - 35, from which 810 subjects have been segmented using the FSL FAST toolset.

For the longitudinal analysis, we found over 4000 MRI scans for a total of 1952 subjects, who have at least 2 visits recorded. Cognitive batteries are notoriously error-prone and subject to tremendous inter-rater variability Mathuranath et al. (2000). Therefore, for the longitudinal cognitive score prediction portion, we found it useful to filter out those subjects whose longitudinal ADAS-Cog data can reasonably be deemed error-prone. Hence in AD subjects, where we do not expect huge *improvements* in cognitive scores with increasing age, we employed an outlier detection routine that removes those subjects whose cognition score increases significantly at any visit from the prior one. The filtered dataset included 1333 scans from 392 subjects with 3 to 7 time points each. Then we used the same split ratio as in the baseline predictions to separate the training set and the set-aside testing set.

### 2.2. Inputs to the model

We used as input to the UNet model: 1) the baseline T1-weighted anatomic MRI scans resolved at 1 mm isotropic voxel resolution; and 2) selected demographics: age, sex, years of education and marriage status. For the categorical variables (marital status and gender) we used one-hot encoding. Our tabular data was then assembled using these demographic variables.

#### 2.2.1. Exclusions

We purposefully omitted regional biomarker features like atrophy, volumetrics, tau and amyloid PET SUVRs – these require advanced processing pipelines. While the longitudinal ADAS-Cog11 data are needed as targets, we did not include any cognitive information at baseline when training the model. Importantly, we also excluded diagnosis, since cognitive performance is strongly governed by the extent of dementia, and diagnosis is in turn offered clinically on the basis of cognitive performance, among other clinical tests.

#### 2.2.2. Primary target 1: Cognitive scores (baseline)

The first outcome is the baseline ADAS-Cog11 score. First introduced by Rosen et al. (1984), the Alzheimer’s Disease Assessment Scale-cognition sub-scale (ADAS-Cog) is widely accepted as a gold standard for assessing the efficacy of anti-dementia treatments. The ADAS-Cog scores are collected through cognition task-based surveys, which are time-consuming, operator-sensitive and burdensome for dementia patients. While other clinical measures like the mini-mental state exam (MMSE) are more common, they are not thought to be true cognitive measures, and display unhelpful bunching at the high end of the scale Cano et al. (2010); Arevalo-Rodriguez et al. (2015). Here we use its summary value, referred to as ADAS-Cog11, of 11 important sub-parts of the test Ito et al. (2011); Samtani et al. (2015). We chose ADAS-Cog11 as our primary research goal because it is predictive of the subjects’ current cognitive status as well as risk of future dementia development Rozzini et al. (2007); Connor and Sabbagh (2008); Kueper et al. (2018).

#### 2.2.3. Primary target 2: Cognitive scores (Longitudinal)

To extend the model to also predict longitudinal ADAS-Cog11, we added a new outcome: the linear slope (change of ADAS-Cog11 per month) beginning at 6 months up to study end, typically 24-48 months. Although condensing the time series to just baseline and slope involves some loss of information, it is an excellent trade-off: a) due to the slow course of disease 2-4 years is a narrow window during which the cognitive trajectory may be reasonably be considered linear, and its slope can be moderately predicted by demographic and clinical data Ito et al. (2011); and b) the slope prediction allows us to trains the same model on subjects with different observation windows, thus increasing addressable sample size. Since there are far more subjects with baseline than longitudinal scores, we trained a separate model for the latter.

#### 2.2.4. Primary target 3: Diagnosis

Incorporating with the ultimate goal of getting clinical advice and treatment of the subjects, we also lay great emphasis on predicting the baseline diagnosis. TODO

#### 2.2.5. Secondary target: Tissue segmentation map

Our secondary target is brain tissue segmentation from baseline MRI. There is no ground truth of tissue segmentation in ADNI, hence we settle for a silver standard obtained from an existing software pipeline called FAST Zhang et al. (2001), a computational tool that segments a 3D image of the brain into 4 tissue types (white matter, gray matter, cerebrospinal fluid and non-brain), whilst also correcting for spatial intensity variations. It is robust and reliable, compared to most finite mixture model methods, which are sensitive to noise Woolrich et al. (2009).

### 2.3. Cross-validation and set-aside data splits

Among these samples we performed a train-test split at a ratio of 9:1. The testing set are set aside and never used in the training and validating processes. On the remaining 90% samples we performed 3-fold cross validation at a ratio of 7:3.

### 2.4. Data augmentation using image pre-processing

Even ADNI, one of the largest standardized imaging datasets of AD, is small in comparison to typical successes of DL. Due to privacy, costs and technical challenges it is usually not possible to achieve much larger samples in medical imaging. Therefore we employed classic data augmentation techniques to artificially generate new training images by applying transformations to existing images; these included: affine image transformations (random crop to 128 x 128 x 128 voxel volumes), elastic transformation to introduce shape variation to the image volumes, and a standard normalization step. The random crop and elastic transformation of the training data increase the generalization ability of the algorithm. These topology-preserving deformations are considered generic and biologically plausible Nalepa et al. (2019). For the validation set, we performed a center crop to 128 x 128 x 128 volumes and standard normalization. Since the validation images serve as the testing ground truth, we use center crop to maintain the same domain as the training set yet capture the same consistent region.

### 2.5. Base CNN Image Model

As the first step, we wish to ascertain whether existing off-the-shelf or “black-box” DL architectures can perform well on the cognition prediction task. Therefore we designed a classical Convolutional Neural Network (CNN) model that takes in the baseline MRI volume and seeks to predict baseline ADAS-Cog11 score. This base CNN network was composed of the following layers: convolutional layer 1, pooling layer 1, convolutional layer 2, pooling layer 2, followed by a 2-layer fully connected Neural Network, and finally the output layer consisting of a single neuron. We adopted the global average pooling layer introduced by popular methods such as ResNet. The output layer utilized a Linear Regression method with a Mean-Squared Error (MSE) loss for the ADAS-Cog-11 score. We experimented first with a convolutional kernel size of 2 × 2 × 2; then by adopting the ideas of Folego et al. Zhang et al. (2001), we enlarged the convolutional kernel size to 3 × 3 × 3 and applied a dropout parameter of 0.5 for regularization. We believe these choices to represent the current state of the art in DL design for the task at hand.

## 3. A Novel Multi-task Network

Since existing ML and DL methods are shown to be sub-optimal for the current task, we developed a bespoke multi-task network that seeks to overcome the twin challenges noted previously: small training sample size and huge dimensionality mismatch. The key idea is that the process of training a model for image segmentation task will produce low-dimensional features that will also be responsive to cognition. The segmentation task will be accomplished using a special type of CNN called a UNet, while cognition will be predicted by a regression block that also incorporates demographic tabular data. The overall multi-task architecture is illustrated in Figure 1. Its components are described below.

### 3.1. Proposed Image Model Architecture: UNet

A convolutional autoencoder is a special type of CNN that is designed to force the image segmentation or regression task to rely on a very low-dimensional latent space of features by bottle-necking the network to go through a series of smaller and smaller layers before expanding back. The UNet is a special type of convolutional autoencoder that alleviates the bottleneck requirement by providing additional connections across downsampled and upsampled feature representations, which improves segmentation accuracy Ronneberger et al. (2015), especially for medical imaging tasks with limited data and the presence of heavy data augmentations Siddique et al. (2021). The architecture consists of: a) a contracting path where the input size is downsampled but the depth of the feature map increases. This allows us to 1) enhance training speed due to the decreasing number of computations and 2) introduce greater generalizability by integrating a small location invariance.

b) an expanding path, where the feature map is up-sampled, that helps regain that precise localization necessary for accurate segmentation. Between the two arms we add “skip” connections where contextual information in the corresponding cropped feature map from the contracting path is concatenated to the expansion layers.

Of key relevance for the cognition task, the lowest-dimensional layer of the UNet (“fully-encoded layer”) is considered to contain contextual information integrated across large brain areas - this is precisely the feature-rich domain that will be expected to carry cognitive information.

The proposed UNet architecture is illustrated in the middle row of Figure 1. The contracting and expanding paths each have four convolution steps. In the contracting path, each layer contains two 3 × 3 × 3 convolutions (with a padding of 1), followed by a rectified linear unit (ReLu), and then a 2 × 2 × 2 max-pooling operation (with strides of two in each dimension). In the expanding path, every layer consists of a 2 × 2 × 2 up-convolution (with strides of two in each dimension), followed by two 3 × 3 × 3 convolutions and a ReLu operation. The last layer consists of a 1 × 1 × 1 convolution that reduces the number of output channels to the number of labels, i.e. 3 labels for our segmentation task.

### 3.2. UNet Loss functions

We use combined objective for training composed of categorical cross-entropy loss and a dice loss. The cross entropy loss is defined as:

The loss function of the UNet segmentation is the cross entropy computed by a pixel-wise soft-max over the final feature map. There are four categories in the ground-truth segmentation map.

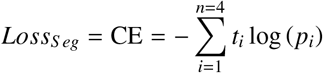

Here *t*_*i*_ stands for the true voxel value, and *p*_*i*_ is the predicted voxel value.

Then, we calculated the weighted Dice coefficient as our metric for evaluation:

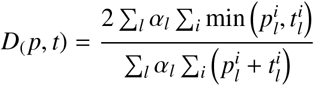

Here 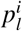 and 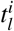 are the set label probability vectors for all voxels for the ground truth and the prediction. The vector *α*_*l*_ denotes the weight of each category.

The loss function for the regression part of the UNet model is the Mean-Squared Error (MSE) loss:

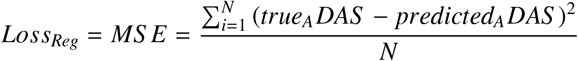

The capital letter *N* denotes the total number of subjects included.

Hence, through integrating the segmentation task and the regression task, we implemented the total loss for the UNet model to be: *L*_*U*_ = *λ*_Cog_ · *L*_Cog_ + *λ*_Seg_ · *L*_Seg_ + *λ*_*α*_ · *L*_*α*_ (*b*) Here, *scale*_*Reg*_ and *scale*_*Reg*_ are the corresponding magnitude of the loss functions, which are divided to balance the power of backward propagation steps, and we initialized the relative importance of the tasks *imp*_*Reg*_ and *imp*_*Seg*_ to be the same.

### 3.3. Ensemble Model Architecture

We wish to enhance the power of our model by adding demographic predictors that are very low-dimensional compared to imaging data - this causes huge dimensionality mismatch whereby the tabular features can easily get lost in translation during training of deep nets. Therefore we selected, after experimenting with various linear and non-linear ML models, a Histogram-based Gradient Boosting regressor (HGB Regressor) from scikit-learn Pedregosa et al. (2011). HGB shows best-tested performance and great stability against over-fitting. We used the Mean Square Error (MSE) as the loss function for the HGB model as well: 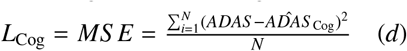

Therefore, we have: Exponential Coeff Loss 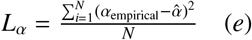

Gamma prior HGB Loss: *y*_emp_ = *y*_true_ + *ϵ*, where *y*_emp_ = *ADAS, y*_true_ is true cognition.

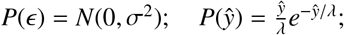

Bayesian posterior 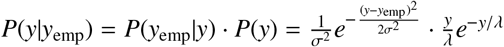

Hence custom HGB Loss

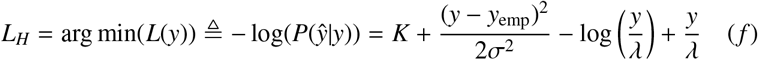

We integrated the regression tasks into an ensemble model by taking the weighted sum of both predictions using their R-squared scores as the indicator of the weights of the corresponding models. We considered an alternate ensemble technique, whereby a multi-input layer was added to the UNet model which brings in the tabular data and made a residual combination in the forward function. We record the R-squared metrics to serve as the weight of each model in the ensemble architecture. Therefore, the loss function for the ensemble model is:

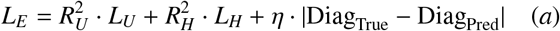

This approach got us comparable results with our best model. Another approach was to train a Gradient Boosting Algorithm along with the UNet and update the loss function of the GB model correspondingly. The performance of these approaches are reported in the Results section.

Table 3 displays the training parameters used during training for the U-Net, segmentation model, the multi-task model, and the ensemble model, respectively.

**Table 1.** Demographic distribution in the dataset: mean (range) over all subjects.

| Characteristic | Training Set | Validation Set | Set-aside Test Set |
| --- | --- | --- | --- |
| Gender (F/M) | 436/554 | 47/65 | 235/253 |
| Marriage (Married/<br>Divorced/Other) | 766/79/145 | 83/7/22 | 360/45/83 |
| Age | 73.2 (55-90.9) | 75.5 (53.4-87.8) | 73.1 (55-91.5) |
| Years of Education | 15.9 (6-20) | 15.9 (12-20) | 16.2 (7-20) |
| MMSE | 26.1 (4-30) | 26.1 (7-30) | 25.5 (0-30) |
| ADAS-Cog11 | 10.9 (1-38) | 10.5 (2-26.7) | 9.5 (1-46.7) |

**Table 2.** *R*^2^ of model prediction on the set-aside testing set, segregated by diagonstic group. While prediction accuracy is comparable across groups, it is lowest for MCI subjects, arguably the most clinically heterogeneous group.

| Diagnostic Group | ADAS-Cog11 range | number of samples in Testing Set | $R^2$ |
| --- | --- | --- | --- |
| Healthy | ADAS ≤ 12 | 320 | 0.8063 |
| MCI | 12 < ADAS ≤ 18 | 67 | 0.7395 |
| Dementia | ADAS > 18 | 101 | 0.7910 |

|  | Segmented Volumes | Loss Function | R2 Confidence Interval on Cog Pred | Average Testing R2 | Diagnosis Accuracy |
| --- | --- | --- | --- | --- | --- |
| MedicalNet50 | True | MSE | 0.73 - 0.82 | 0.6738 (0.6553 - 0.7421) | 0.9251 |
| MedicalNet50 | True | Gamma Loss | 0.68 - 0.84 | 0.7113 (0.6603 - 0.7548) | 0.9415 |
| MedicalNet50 | False | Gamma Loss | 0.54 - 0.72 | 0.6126 (0.5771 - 0.6424) | 0.8949 |
| Multi-task UNet | True | MSE | 0.63 - 0.77 | 0.6665 (0.6210 - 0.7095) | 0.9090 |
| Multi-task UNet | True | Gamma Loss | 0.70 - 0.81 | 0.6919 (0.6748 - 0.7254) | 0.9437 |

**Table 3.**
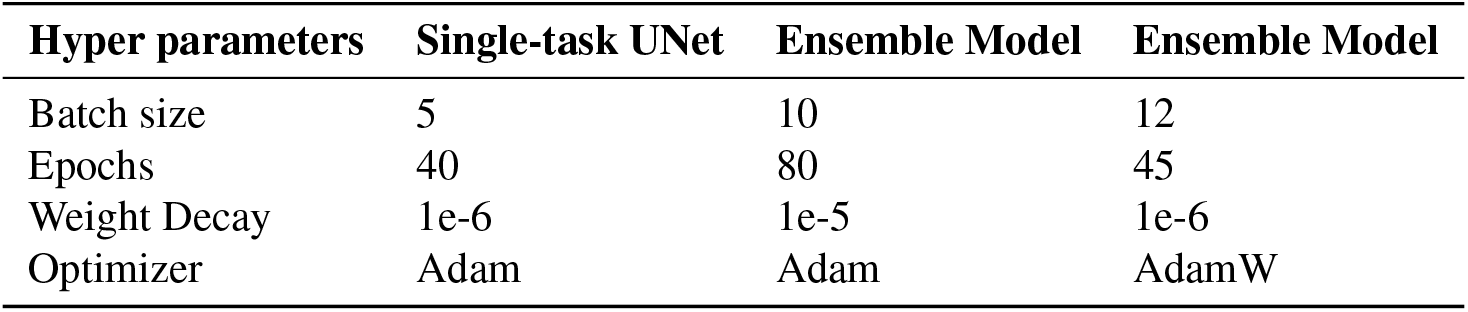
Training Parameters for Each Model.

### 3.4. Feature Map Analysis

To gain insight into how UNet-derived features enable enhanced prediction of ADAS-Cog scores, we utilize the local receptive field analysis method developed recently by Rajagopal et al. (2021) to visualize and quantify how the feature-space correlates with cognitive score, patient pathology, and different MRI phenotypes at various layers of the UNet. Specifically, this is achieved by extracting the neural trace derived from the training data (MRI exams) of the UNet model, applying the local receptive field analysis method to extract correspondences between quantized features across layers of the model, and visualizing these correspondences as point-clouds and dense volumetric slices with coloring representing different semantic groups (feature phenotype, cognitive score, etc.).

### 3.5. Longitudinal Cognition Prediction

To achieve the clinically important objective of forecasting future cognitive scores from baseline MRI, we built a separate model mirroring the above architecture, with two differences:

a. Instead of baseline cognition, the outcome measure is the cognitive slope (change per month), computed for each subject by fitting a straight line starting from baseline cognition (zero intercept). Slope outcome can handle different visiting timelines of subjects without retraining.
b. We added a new input feature from the segmentation map produced by the UNet: z-score of the total volume of each segmented tissue class compared to healthy controls. This exploits the well-known dependence of cognitive decline on baseline atrophy Rusinek et al. (2003); Kramer et al. (2007); Whitwell et al. (2008).

Although it is theoretically possible to add the slope outcome in the same multi-task architecture, a separate model is more appropriate here since it needs to be trained on a different, smaller, sample size, while leveraging segmentation output that was not necessary for the baseline model.

## 4. Results

### 4.1. Segmentation Performance

As shown in Figure 2, the UNet architecture was able to generate segmentation results visually indistinguishable from ground truth. The Dice similarity coefficient (DSC) was used as the validation metric of reproducibility of manual segmentations and spatial overlap accuracy. Our best model achieved validation DICE coefficients of 0.9978, 0.9635, 0.9513, and 0.9725 for background, gray matter, white matter, and cerebrospinal fluid, respectively. The overall DICE coefficients, listed in Table **??**, compare favorably with most cutting-edge segmentation algorithms (see Discussion).

**Fig. 2.**
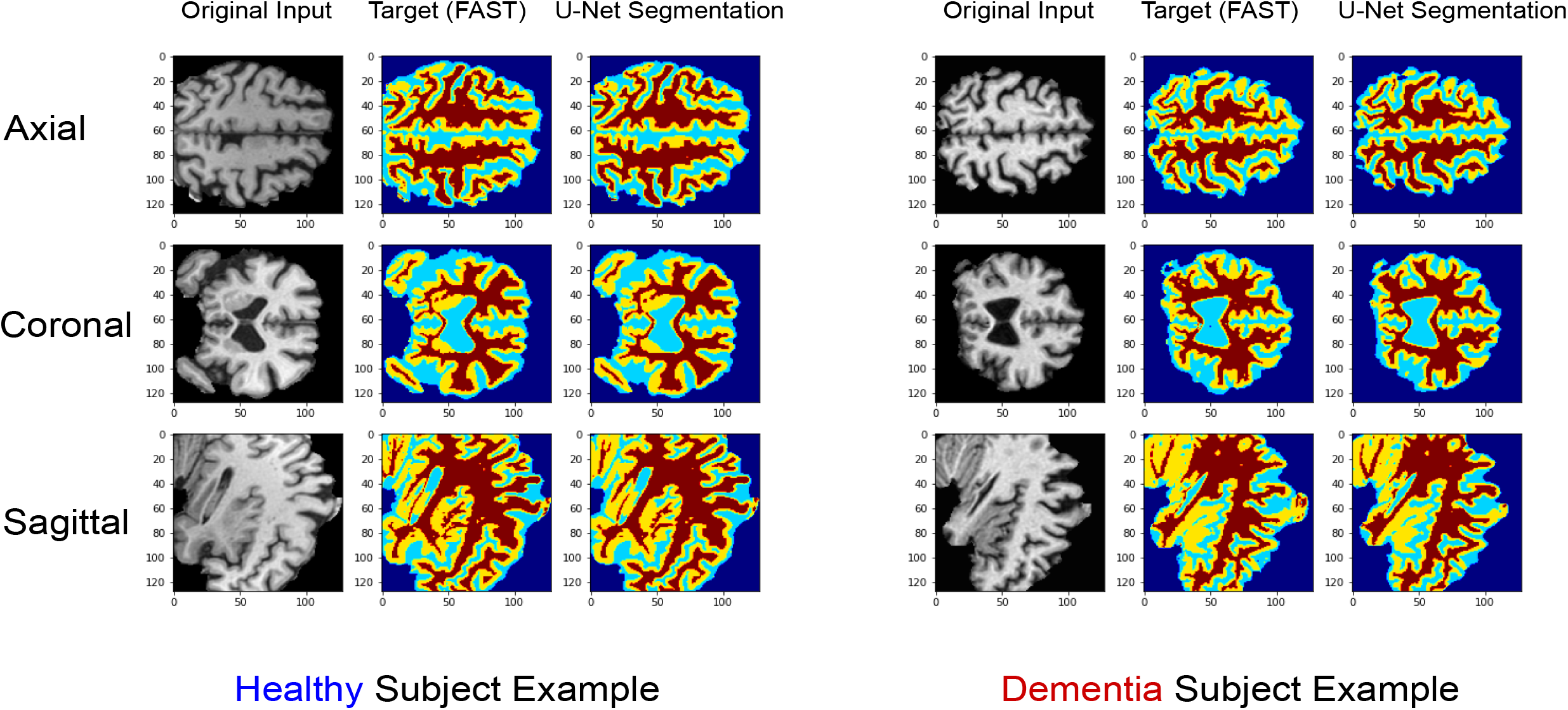
Two representative examples of our image segmenta. tion outputs. The raw MRI image is shown alongside the FAST-generated segmentation map which constitutes our silver-standard target, and the UNet-generated segmentation result. Three orthogonal slice orientations are shows: axial, coronal and sagittal. The subject on the left belongs to CN group, while the right subject belongs to Dementia group.

**Fig. 3.**
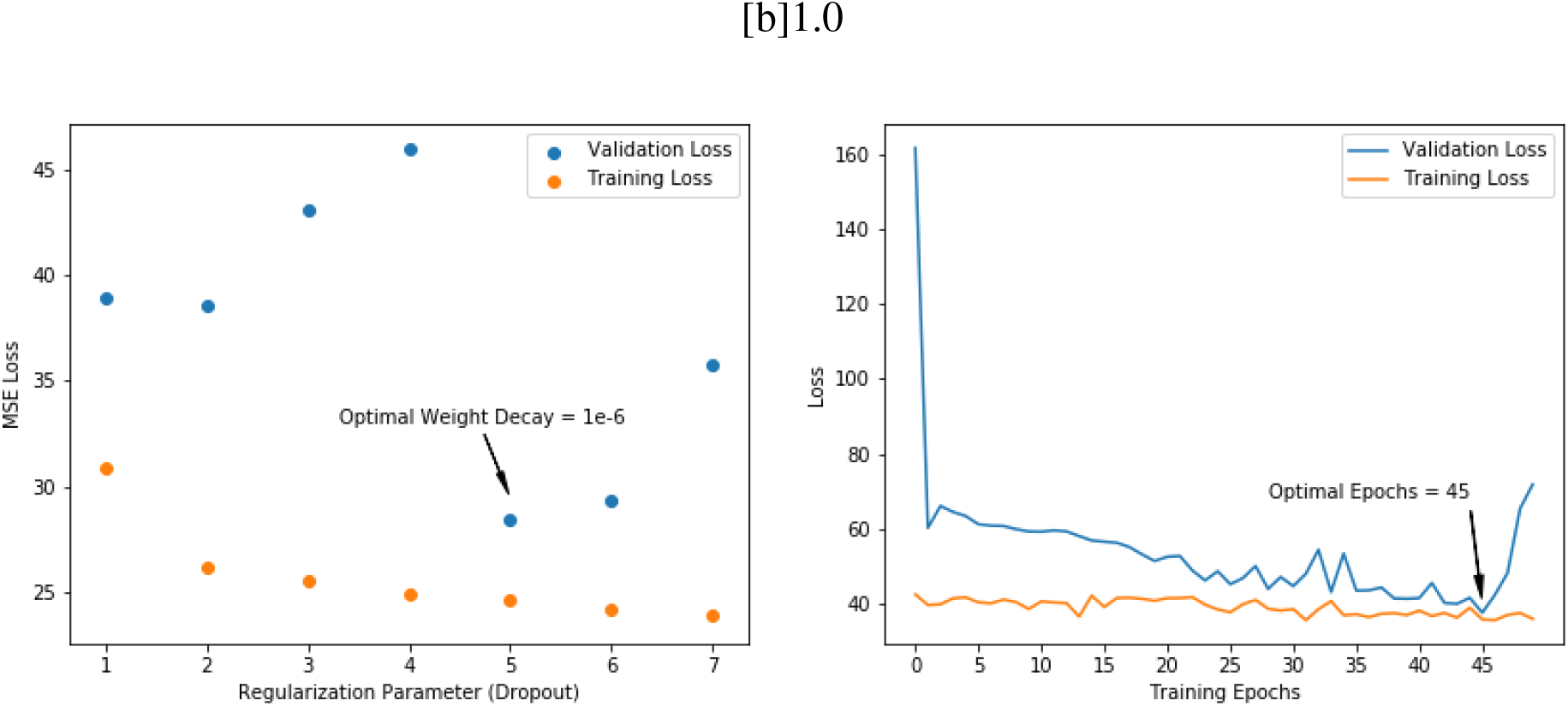

**Fig. 4.**
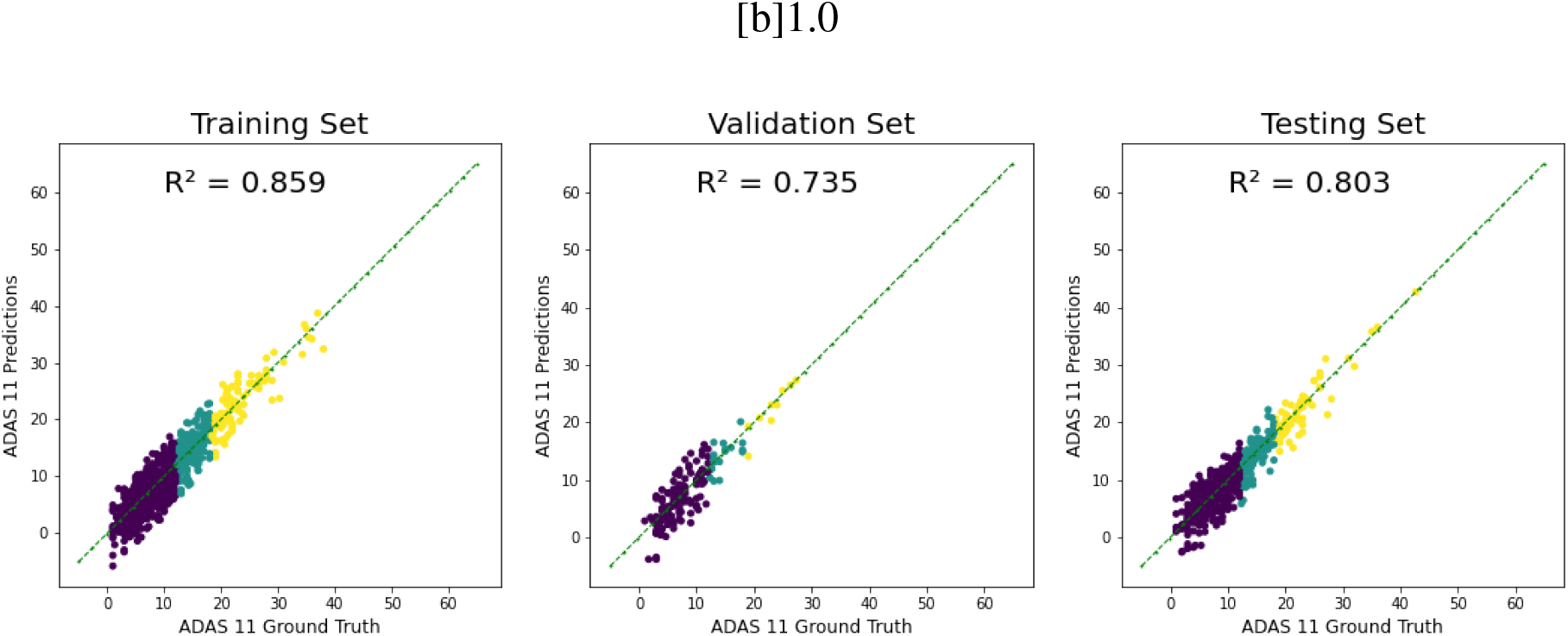

### 4.2. Regression Performance

Table **??** summarizes the experimental results for the baseline cognition task of all models we tested. All *R*^2^ reported in this section pertain to the set-aside testing set, using the training/validation/set-aside split described below. The HGB regressor alone produces *R*^2^ over 0.5, revealing that the tabular demographic data are decent predictors of baseline cognitive score. Our implementation of the single-task CNN model achieved mediocre *R*^2^ on the set-aside testing set, lower than HGB, supporting our intuition that a more complicated, custom architecture is necessary. The Multi-task UNet model gives the best, simultaneously achieving accurate segmentation and lower training loss and higher *R*^2^ on ADAS-Cog regression task than both the tabular model and the single-task CNN model. While the UNet’s regression task is already accurate (*R*^2^ = 0.59), by concatenating it into the ensemble method we achieve even higher accuracy, as described below.

We experimented on several regression models that take as input the fully-encoded layer of the UNet and predicts cognition. Among these PyTorch’s MLP (2 NN Linear layers) outperformed others. We also evaluated Random Forest, XGB, and kernel-based Support Vector Machine as non-linear substitutes for the MLP, mainly because we found that the intermediate outputs from each convolution block had strong nonlinearity (see section 5.2). However, the neural network’s MLP benefited from its ability to generate a large number of parameters and to absorb the nonlinearity using ReLU activation functions.

#### Ensemble model

Our first ensemble attempt was to design a 2-layer MLP that took in both the tabular demographic data and the UNet bottom-layer latent vectors. This gave us MSE of 34.13 plus an *R*^2^ = 0.68. Next, we trained a CNN for the prediction task alone using ADNI Image data. Those results are reported in Table **??**. Then, inspired by the idea of ResNet He et al. (2015), we developed a specific *R*^2^ ensemble model, assuming that *R*^2^ of each separate model measures the amount of interpretability it adds to the whole task. We assigned the original *R*^2^ of both the UNet segmentation task and the GBR model as their parameter weights, and stacked these two models to form the R-squared Ensemble Model.

#### Hyper-parameter tuning and cross-validation of ensemble model

Figure 5(a) shows the effect of hyper-parameter selection in the final ensemble model based on the performance of 10-fold cross-validation. First, we fixed the dropout levels of the model at 1e-6, 1e-7, and 1e-8, and printed out the validation MSE loss versus the training epochs. The ensemble model reached its optimal validation results on 45 training epochs in all three dropout levels. Then, we fixed the training epoch to be 45 and found that a value of 1e-6 gives the best dropout regularization parameter for our U-Net-based multi-task model. Figure 5(b) shows the *R*^2^ performance on the baseline cognition task for the three sets: one group of training set (*R*^2^ = 0.86), the respective cross-validation set (*R*^2^ = 0.74) and set-aside testing set (*R*^2^ = 0.80). While all three sets produced excellent accuracy, as expected the training set achieves the highest, but the out-of-sample set achieves nearly the same accuracy, higher than the validation set. Through sanity check, we confirmed that this is majorly because the testing set has a similar underlying distribution with the training set.

**Fig. 5.** Hyper-parameter Tuning and Corresponding Prediction Results. (a) Left: Training/validation performance at different choices of weight decay. The optimal decay level is 1e-6. (a) Right: Training/validation performance at fixed weight decay (1e-6) and increasing training epochs. We chose 45 epochs, which give the lowest validation loss. (b) Scatter plots of performance of the ensemble model in predicting baseline ADAS-Cog11, separately for subjects within the in-sample training set, validation set and out-of-sample test set. All sets give excellent correlation with real data.

All results shown in Table **??** pertain to the set-aside test set. The best-performing ensemble model is listed in the last row (*R*^2^ = 0.80). Interestingly, model performance is excellent across all diagnoses; Table 2 shows *R*^2^ of model prediction on the set-aside testing set, segregated by diagonstic group. While prediction accuracy is comparable across groups, as expected it is lowest for MCI subjects, arguably the most clinically heterogeneous group.

### 4.3. Prediction of longitudinal cognition

Figure 6A plots empirical versus predicted ADAS-Cog11 slope from the longitudinal trained model. The top panel corresponds to the case where the model is trained without the additional segmentation class volumes as inputs; this approach achieved *R*^2^ = 0.69. Then, introducing the tissue-class segmentation volumes computed from the UNet’s baseline segmentation image output as additional inputs into the linear layer with a weight of 0.3 (empirically chosen), we retrained our best ensemble model. This tissue-volume-enhanced approach was able to significantly improve performance of slope prediction, achieving *R*^2^ = 0.77. This result, shown in the bottom panel, represents our best result on the slope task.

**Fig. 6.**
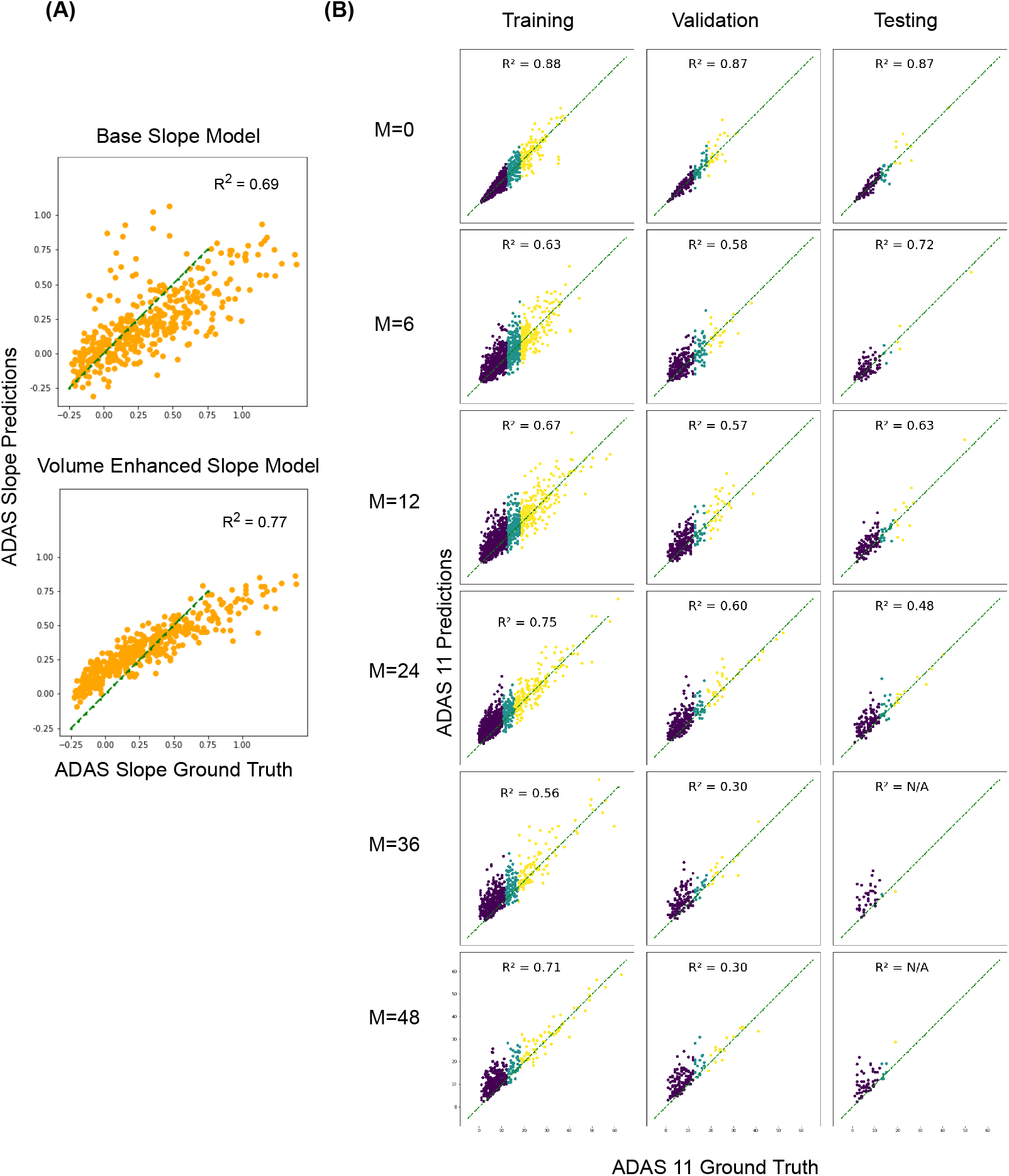
Prediction of longitudinal decline in cognition. A. Scatter plots of ADAS-Cog11 slope (change per month), empirical vs predicted. The top plot shows the results from the base longitudinal model, which simply changes the prediction target from baseline ADAS-Cog11 to its slope. The bottom plot shows results of the volume-enhanced model, obtained after re-training the UNet with an additional input: the tissue class volumes from the segmentation output. B. Scatter plots of ADAS-Cog11 prediction for individual longitudinal visits, reported separately for 6,12,24,36 and 48 months. The x-axis of each subplot is the empirical ADAS-Cog11, and y-axis is the predicted value. Subjects are color coded by diagnosis: blue=control, cyan=MCI, and yellow=AD.

The next group of scatter plots in Figure 6B reports the predictions of cognitive scores segregated by month of visit. The x-axis of each subplot is the empirical ADAS-Cog11, and y-axis is the predicted value. Clearly, the predictions of ADAS-Cog11 are highly accurate for up to 24 months, thereafter they begin to decline in accuracy. However, even up to 48 months the training and in-sample validation sets exhibit excellent accuracy. It is only in the out-of-sample test set that the accuracy plummets beyond 24 months; this is likely due to the exceedingly small sample sizes pertaining to this group at such long follow up visits.

### 4.4. Feature space visualization

Feature analysis using a novel visualization approach Rajagopal et al. (2021) for the 3D UNet, shown in Figure 7A, highlights alignment between the features extracted by the segmentation model and the ADAS-Cog scores, providing further confidence in the success of our hybrid ML. Figure 7B solidifies this point by demonstrating semantic segmentation of brain tissue at intermediate layers of the network, despite not being trained for this task. In particular, these features appear to vary as a function of cognitive score on validation and test patients, indicating that the UNet-extracted features may serve as generalizable biomarkers for AD and other neurological diseases.

**Fig. 7.**
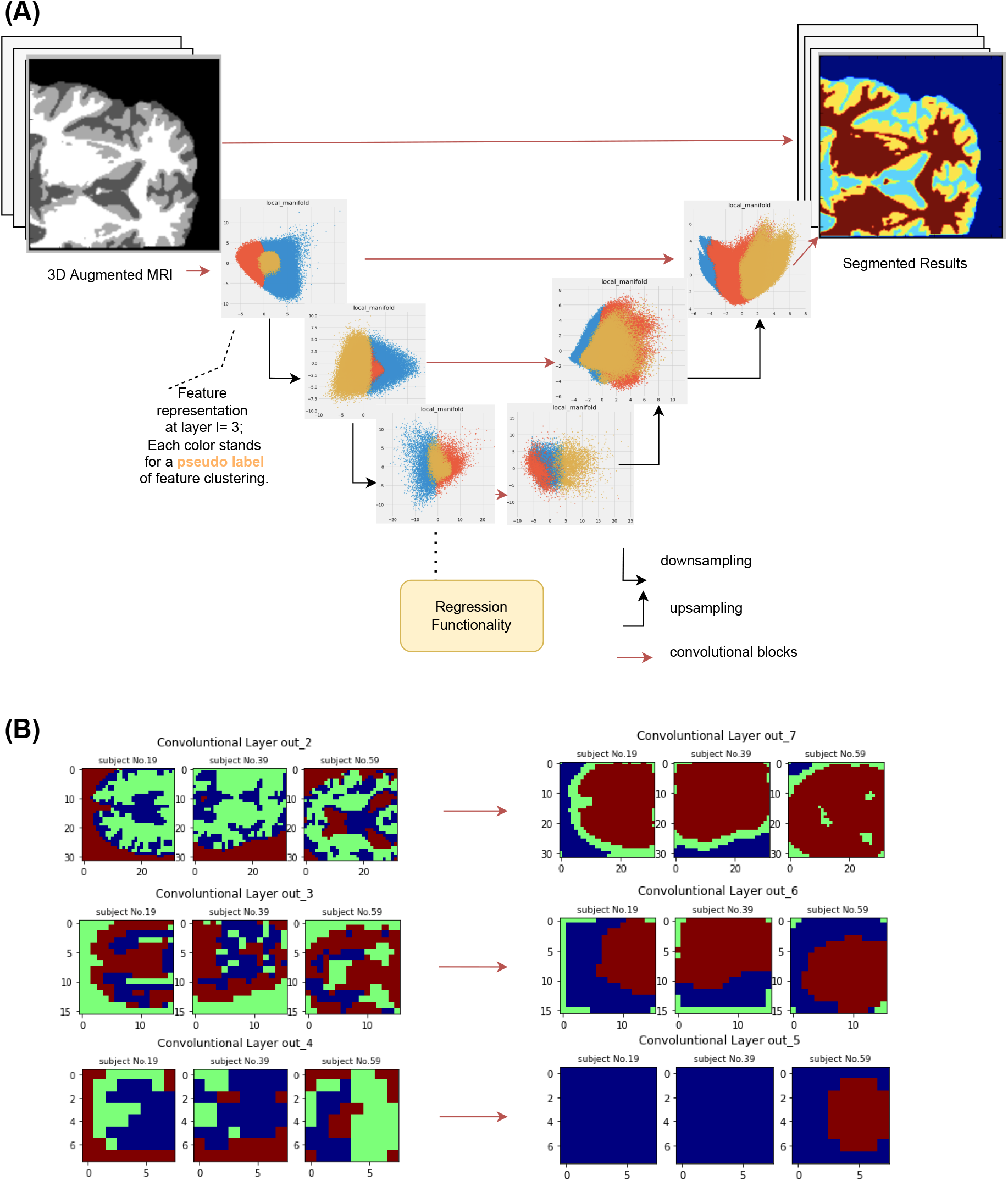
A. Feature Map Visualizations. Local receptive field feature representation of our 3D UNet trained on the 4-category segmentation of the Structural MRI, depicting extraction of intermediate feature manifolds with corresponding pseudo-labels indicated by color. **B. Voxel-level Visualizations. Here we selected three subjects’ clustered slices from each layer. Each color stands for a pseudo-label, and contribute differently to the regression scores. Layer out4 is the last layer before the regression block, for which reason it concentrates the most informative features for the ADAS prediction. As the UNet switch to upsampling function starting from out5, the attention of the pseudo-labels move to re-generating the segmented images**.

## 5 Discussion

The goal of this study was to design and test a novel neural network model for two tasks relevant in Alzheimer’s disease research and clinical practice: how to quickly perform image segmentation into gray, white and CSF tissue classes in the brain; and how to predict a patient’s cognitive score from their MR image and common demographic and clinical information typically collected during standard-of-care exams.

### 5.1. Summary of key findings

Due to the challenges inherent in the dimensionality mismatch between images and cognitive scores, it was found that off-the-shelf black-box neural network models fail to achieve reasonable results - this was confirmed by our implementation of the baseline single-shot CNN model that achieved poor performance on the cognitive prediction task. Therefore we designed a novel multi-task model involving a UNet that exploits the image features learned during the segmentation task as input features for the cognitive task. At the same time, the model was designed to accommodate scalar or tabular demographic data – age, sex, year of education, marriage status, etc – that are also known to be strong predictors of cognitive performance. The challenge inherent in combining voxel-level, tabular and scalar data was ameliorated via a bespoke multi-task design. The multi-task model simultaneously performed segmentation of white matter, gray matter, and cerebrospinal fluid on the MRI input and regression to predict the ADAS-Cog11 score. Both tasks were accomplished to a best-in-class level of accuracy. In this, it exploited the well-known literature highlighting the independent relationships between gray and white matter volumes and dementia severity, in addition to other more direct MRI image features.

Our final multi-task model, combining 3D UNet and Gradient Boosting Trees, comfortably out-performed other tested models for prediction of cognitive scores. Prior DL applications in the neuroimaging space have been rightly criticised for overfitting, poor cross-validation, lack of biological interpretability and the use of small sample sizes Sui et al. (2020). We were able to address and overcome these concerns by utilizing a rigorous training / validation / set-aside testing regime, thereby mitigating the risks of overfitting while employing one of the largest sample sizes available in the field. In contrast to many ML models that lack interpretability, we leveraged our prior theoretical work Rajagopal et al. (2021) to obtain visualizations of the operative features’ receptive fields and their clustering in feature space.

Of particular importance is our demonstration that the presented model can correctly predict future levels of cognition, in addition to accurately predicting baseline tissue segmentation and cognition. Extending our approach to full longitudinal cognitive prediction required re-utilising the tissue-wise volumes calculated from the segmentation output - an excellent example of biologically motivated data reuse and transfer learning. As a result our model was able to give excellent predictions of future cognition, up to 48 months out, using baseline imaging and demographic data. The resulting model may therefore become useful as a fast, easily implementable and highly accurate tool that may be used in most common clinical settings, without requiring specialized tools or expensive PET scans, genetic analysis or fluid proteomics.

### 5.2. MRI biomarkers to cognition: existing methods and challenges

Given the highly complex nature of the brain, it may be asked, why an approach that primarily uses structural MRI can predict cognition at all, since cognition is considered an emergent property that may only be loosely related to specific brain structures and other morphological features. Indeed, it would be challenging to reproduce our results on similar high-level emergent properties like intelligence. Here, we are aided by the remarkable effect that neurodegeneration has on the brain’s morphology. Thus, by training our model on a sufficiently diverse and large group of AD-spectrum subjects at various stages of neurodegeneration, we can learn the complex but deterministic relationship between morphology and cognition. Our approach thus implicitly exploits the strong temporal ordering of multiple AD biomarkers Jack and Holtzman (2013): genetic predisposition causes tau and amyloid deposits, which subsequently spread into wider circuits and cause atrophy and structural changes that then impact cognitive and clinical function.

The association between longitudinal changes in MRI and cognitive decline is well-studied, especially their derivative features like regional volumes and thickness. Several brain regions critical for cognition, e.g. hippocampus, become smaller with age, and structural brain changes in healthy elderly at baseline predicted a more rapid cognitive decline Rusinek et al. (2003) and higher rates of conversion to dementia. Changes in hippocampal volumes are associated with decline in memory scores while cortical volume changes are associated with decline of executive function in normally aging subjects Kramer et al. (2007). MRI-derived temporal atrophy was higher in MCI subjects who progressed to AD compared to those who were stable Whitwell et al. (2008). However, these associations are not always significant, with conflicting data; e.g. an earlier study found no significant relationship between hippocampal atrophy and memory decline Ylikoski et al. (2000). Hence a clear understanding of the mechanisms underlying cognitive aging remain unclear Kramer et al. (2007).

In comparison to prior volumetric analyses, the present approach presents clear benefits, as it circumvents the need to employ highly specialized, manual, time-consuming and computationally demanding MRI volumetry software. Further, volumetry methods have been demonstrated in longitudinal settings, employing changes in MRI-derived volumes over time, and their ability to operate with only baseline MRI is much poorer Lye et al. (2004); Van Petten (2004). An excellent meta-analysis of the challenges of prediction of cognition from MRI biomarkers Sui et al. (2020) shows that prediction accuracy ranges from 0.2 to 0.8 in the vast majority of studies, with a mean around 0.5. These numbers are much lower than achieved by the current approach that allows the network to learn the right representation of morphological structures in a parsimonious manner without relying exclusively on specific brain regions or shapes.

### 5.3. Comparison to Prior Deep Learning Work

Many prior studies have effectively explored the use of machine learning, deep learning and CNNs to achieve accurate diagnosis from baseline imaging data in AD Suk et al. (2014); Pachauri et al. (2011) and predicting conversion to dementia Davatzikos et al. (2009, 2011); Mattsson et al. (2014); Pachauri et al. (2011); Risacher et al. (2009). Of these, 6 DL studies met the PRISMA guidelines between January 2017 and August 2021, reported in the survey Wassan et al. (2021). Automated diagnosis on a small sample of 34 patients achieved 95.59% diagnosing accuracy using MRI biomarkers Qiao et al. (2018). Others have applied 3D DNN on PET images to diagnose AD (84.2% accuracy) Choi and Jin (2018a), or 2D CNN (AUC over 98%) Ding et al. (2019b). A 3D CNN using rs-fMRI reported accuracy over 90% Qureshi et al. (2019). A very large ResNet model trained on MRI pooled from more than 217 sites/scanners gave classification accuracy of 91.3% Lu et al. (2021). A multi-modality CNN trained on 2861 MRIs achieved AUC of 92.01% Huang et al. (2019). An inception V3 network trained on PET performed better than radiology readers, in predicting MCI converters Ding et al. (2019a). A 3D CNN was employed to predict conversion from MCI to AD using PET scans with an accuracy of 84.2% Choi and Jin (2018b), without the need for morphometric features.

Despite these successes, there is a relative dearth of ML/DL approaches for the more interesting task of predicting a subject’s cognitive score from imaging. We list below a few such studies. However, we note that to our knowledge, *none of them have demonstrated the ability to predict cognition into the future using baseline imaging*. As noted in a recent review, compared with other fields, the application of DL to predictive modeling in neuroimaging is relatively modest, due to lack of interpretability and the need for extensive amounts of data Sui et al. (2020).

Back in 2010, Relevance vector regression, a sparse kernel method, was proposed for the prediction of clinical ratings and ADAS-Cog from morphometric analysis of MRI Stonnington et al. (2010), reporting *R* = 0.57. Considering this was work 10 years ago, this is excellent performance, albeit without current standards of cross-validation. Further, this approach is limited by intensive post-processing pipelines for computing deformation fields, which can slow down the training process. More recently, a multi-task dictionary learning CNN model pre-trained on the rich ImageNet data set Zhang et al. (2017) was able to predict cognitive scores, after transfer-learning to take the baseline volumetric and thickness features from hippocampus, ventricles and cortex as input. The model produced combined *R* = 0.862, roughly equivalent to our results. Of course, this method required not only vastly larger training data and training times but morphometric processing pipelines. Notably, their approach did not include a set-aside testing set, hence its generalizability and potential risk of overfitting remain unaddressed.

### 5.4. Strong translational and clinical potential

The ability to predict a patient’s cognitive score from imaging opens up many meaningful opportunities for clinical utility and intervention. It may be emphasized, neurologists base their clinical diagnosis primarily on cognitive status combined with subjective clinical assessment. The ability to relate imaging to cognition may therefore greatly assist the neurologist in solidifying diagnostic and prognostic decision-making. We reported excellent prediction of not only baseline but also *future* cognitive scores, a key element of computational and personalized staging, prognosis and early detection. The unique strength of our study is that it uses Deep Learning on baseline MRI - not requiring longitudinal imaging and circumventing prior reliance on specialized MRI morphometric software or operator-selected regional atrophy measurements. Hence our approach is more liable to be translated and adopted in community clinics that do not possess those highly specialized skill sets. The approach presented here is not limited to AD and may prove relevant for clinical trials of disease-modifying drugs, and other neurodegenerative diseases like Parkinson’s, ALS, and Huntington’s disease.

### 5.5. Limitations and Next Steps

The current study focuses exclusively on MRI and demographics as input features. The prediction of cognitive decline will certainly be improved by additional imaging modalities and other data, especially longitudinal ones. There is a possibility of combining the present model with biophysical models of atrophy progression using e.g. our laboratory’s prior Network Diffusion Model Raj et al. (2012). Hence, a model that could learn and combine the temporal and regional associations between all available biomarkers from multimodal imaging (*Aβ*- and Tau-PET, MRI-derived atrophy, and FDG-PET), could further improve its performance and enable complete computational tracking and prediction of disease progression. Although MRI remains the most accessible diagnostic tool, genetic Xu et al. (2022), blood and CSF protein levels are emerging as exciting new biomarkers that can aid the prediction of cognition Dickerson and Wolk (2013).

We also note the limitation of pure ML approaches in interpretability and understanding of disease processes. We partially alleviated this by performing thorough feature space analysis that clearly links image-level features with group-level clustering of phenotypes. Finally, while DL methods are very effective at learning associations, they typically require very large training sets numbering in the millions – those numbers are clinically unfeasible. We employed several strategies to overcome the sample size issue, yet future studies will benefit from more training data.

---

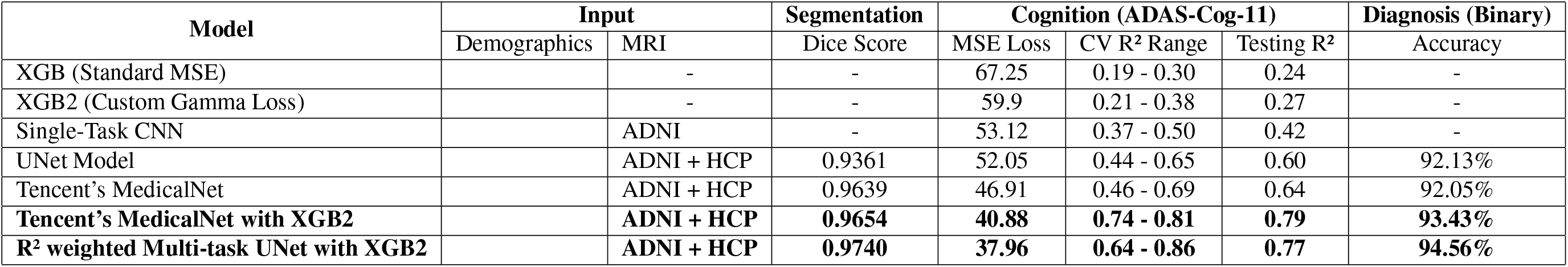

## Notes

### Competing Interest Statement

The authors have declared no competing interest.

### Summary of Updates

This version of the manuscript has been revised to update the following: 1. More training data are involved in the cohort. We enlarged the dataset from only ADNI1,2 to full ADNI plus HCP-YA images. 2. We experimented with more benchmark models more rigorously, and provided a customized loss function for the deep learning part. 3. We added more QoL changes to the model including RFA analysis and so on. We believe this version is a better expression of our research results and study progress.

